# A phenotypic screening platform for chronic pain therapeutics using all-optical electrophysiology

**DOI:** 10.1101/2022.12.12.520139

**Authors:** Pin W. Liu, Hongkang Zhang, Christopher A. Werley, Monika Pichler, Steve Ryan, Caitlin Lewarch, Jane Jacques, Jennifer Grooms, John Ferrante, Guangde Li, Dawei Zhang, Nate Bremmer, Adam Barnett, Romina Chantre, Amy E. Elder, Adam E. Cohen, Luis A. Williams, Graham T. Dempsey, Owen B. McManus

**Author notes:** Pin W. Liu, Hongkang Zhang and Christopher A. Werley contributed equally to this work. Corresponding author: Owen McManus, email address, 179 Sidney St., Cambridge, MA 02139.

## Abstract

Chronic pain associated with osteoarthritis (OA) remains an intractable problem with few effective treatment options. New approaches are needed to model the disease biology and to drive discovery of therapeutics. Here, we present an *in vitro* model of OA pain, where dorsal root ganglion (DRG) sensory neurons were sensitized by a defined mixture of disease-relevant inflammatory mediators, here called Sensitizing PAin Reagent Composition or *SPARC*. OA-SPARC components showed synergistic or additive effects when applied in combination and induced pain phenotypes *in vivo*. To measure the effect of OA-SPARC on neural firing in a scalable format for drug discovery, we used a custom system for high throughput all-optical electrophysiology. This system enabled light-based membrane voltage recordings from hundreds of neurons in parallel with single cell resolution and a throughput of up to 500,000 neurons per day, with patch clamp-like single action potential resolution. A computational framework was developed to construct a multiparameter OA-SPARC neuronal phenotype and to quantitatively assess phenotype reversal by candidate pharmacology with different mechanisms of action. We screened ~3000 approved drugs and mechanistically focused compounds, yielding data from over 1.2 million individual neurons with detailed assessment of both functional OA-SPARC phenotype rescue and orthogonal “off-target” effects. Analysis of confirmed hits revealed diverse potential analgesic mechanisms including well-known ion channel modulators as well as less characterized mechanisms including MEK inhibitors and tyrosine kinase modulators, providing validation of the platform for pain drug discovery.

## Introduction

Pain remains the primary reason patients seek medical care[88]. Chronic pain is difficult to treat, leading to significant morbidity, loss of productivity and lower quality of life[19,49,79]. Among chronic pain indications, osteoarthritis (OA) is particularly prevalent[37,64,48,52], with >30 million OA patients in the US, representing about one third of the total chronic pain population[81]. OA and other forms of chronic pain are widely treated with opioids, but mechanism-based adverse effects, tolerance, and dependency limit long term use, leading to a major health crisis in the US[31]. Gabapentinoids carbamazepine and nonsteroidal anti-inflammatory agents provide relief for only a subset of patients[34]. Despite the clear unmet medical need and significant research activity, few effective non-opioid drugs for OA pain have appeared in the past two decades.

A major factor limiting development of pain therapeutics is lack of scalable, translatable models of sensory neuron function, which is essential for transmission of pain signals from the periphery to the brain[39]. Traditional approaches rely on target-based high-throughput screens, often under conditions that do not mimic physiological environments, followed by *in vivo* evaluation using animal pain models that often do not translate clinically [30]. Assays based on cell-based models have traditionally suffered from severe tradeoffs between throughput and information content.

Here we address this need through development of a high-throughput, all-optical electrophysiology platform focusing on an OA “*pain-in-a-dish*” model in rat dorsal root ganglion (DRG) neurons sensitized with OA-SPARC (OA-derived Sensitizing PAin Reagent Composition). The OA-SPARC is an optimized formulation of inflammatory mediators found at elevated levels in synovial fluid of arthritic joints of patients[66]. To enable high-throughput screening, we used simultaneous optical stimulation and optical recording of neuronal action potentials (APs) with genetically encoded actuators and voltage indicators [38]. We previously developed the Firefly, a multi-well plate compatible, widefield microscope for optogenetic recordings from hundreds of neurons simultaneously with millisecond temporal resolution and single cell spatial resolution[67]. This system maintains the rich information of manual patch clamp voltage measurements, with >30,000-fold higher throughput. Using diverse stimulation patterns, we extract >500 electrophysiological features per neuron. OA-SPARC application drives a strong, multiparameter hyperexcitability phenotype in cultured DRG neurons and causes pain *in vivo* when injected into a rat hindpaw.

We pharmacologically validated the model by showing phenotype reversal with clinical analgesics and expected responses to potent and selective compounds with diverse targets. We performed a phenotypic screen of ~3000 FDA-approved and mechanistically focused compounds to identify potential analgesic targets. Analysis of confirmed hits revealed known analgesic mechanisms (e.g. ion channel modulators), and less well characterized mechanisms (e.g. MEK inhibitors). Combining high throughput optical electrophysiology with a rationally designed pain sensitization model enabled detailed phenotypic measurements for drug discovery in pain-relevant models of sensory neuron function. This platform promises to be more informative and relevant to human physiology than heterologous expression models, and faster, cheaper and more quantitative than *in vivo* models.

## Results

### High-throughput, single-cell measurements of sensory neuron excitability for assessing candidate pharmacology

We previously built an integrated technology platform combining cell-based models, custom instrumentation[38] for optogenetic recordings, and computational tools for high-throughput measurements of neuronal excitability[67,97,96,98] and synaptic transmission[8]. The engineered channelrhodopsin CheRiff enables neuronal stimulation with blue light and the engineered voltage-sensitive protein QuasAr enables high-speed fluorescence recordings of changes in membrane potential with red light[106]. Our wide field-of-view (FOV) *Firefly* microscope (0.8 × 4 mm) can stimulate any subset of neurons with a fully configurable optical pattern using a digital micromirror display and simultaneously record voltage from the entire FOV with single-cell spatial resolution, millisecond temporal resolution and a signal-to-noise ratio (SNR, spike height:baseline noise) of approximately 9.5 (95% CI: 4.58-20.3). Custom software and analysis tools handle the high-speed voltage imaging data. The Firefly system has been applied to many types of rodent primary neurons and human induced pluripotent stem (hiPS) cell-derived neurons[44,96,98].

Here we extended our approach to *in vitro* cultures of rodent DRG sensory neurons. Rat DRG neurons were cultured in 96-well plates, and optogenetic constructs were introduced through lentiviral delivery. Movies were then recorded serially, well-by-well on the *Firefly* microscope. Neurons within each FOV were interrogated with a varied stimulus pattern using blue light (470 nm) designed to probe a broad range of neurophysiological behaviors. We established an automated analysis workflow. After recording from each plate, the typically 100s of Gigabytes of data were transferred to Amazon Web Services (AWS) and hundreds of processors analyzed the movies in parallel. An arbitrarily large dataset could be analyzed in ~1 hour by scaling the number of processors. Results were downloaded into an internal database for further analysis.

We validated the platform using an established mixture of soluble mediators that occur in inflammatory pain[4], here called ‘Inflam-SPARC’. Figure 1 shows optical voltage recordings from over 11,000 individual neurons obtained from 1/3 of a 96-well plate. Two conditions are shown: 1) neurons treated with Inflam-SPARC[89] and 2) vehicle control-treated neurons. A FOV of ~100 DRG sensory neurons in the presence of supporting cells (Fig. 1A) was stimulated with a protocol that included short (100 ms) and long (500 ms) depolarization steps with various blue light intensities and a linear depolarizing ramp at the end (Fig. 1B, see Methods for detailed protocol).

**Figure 1.**
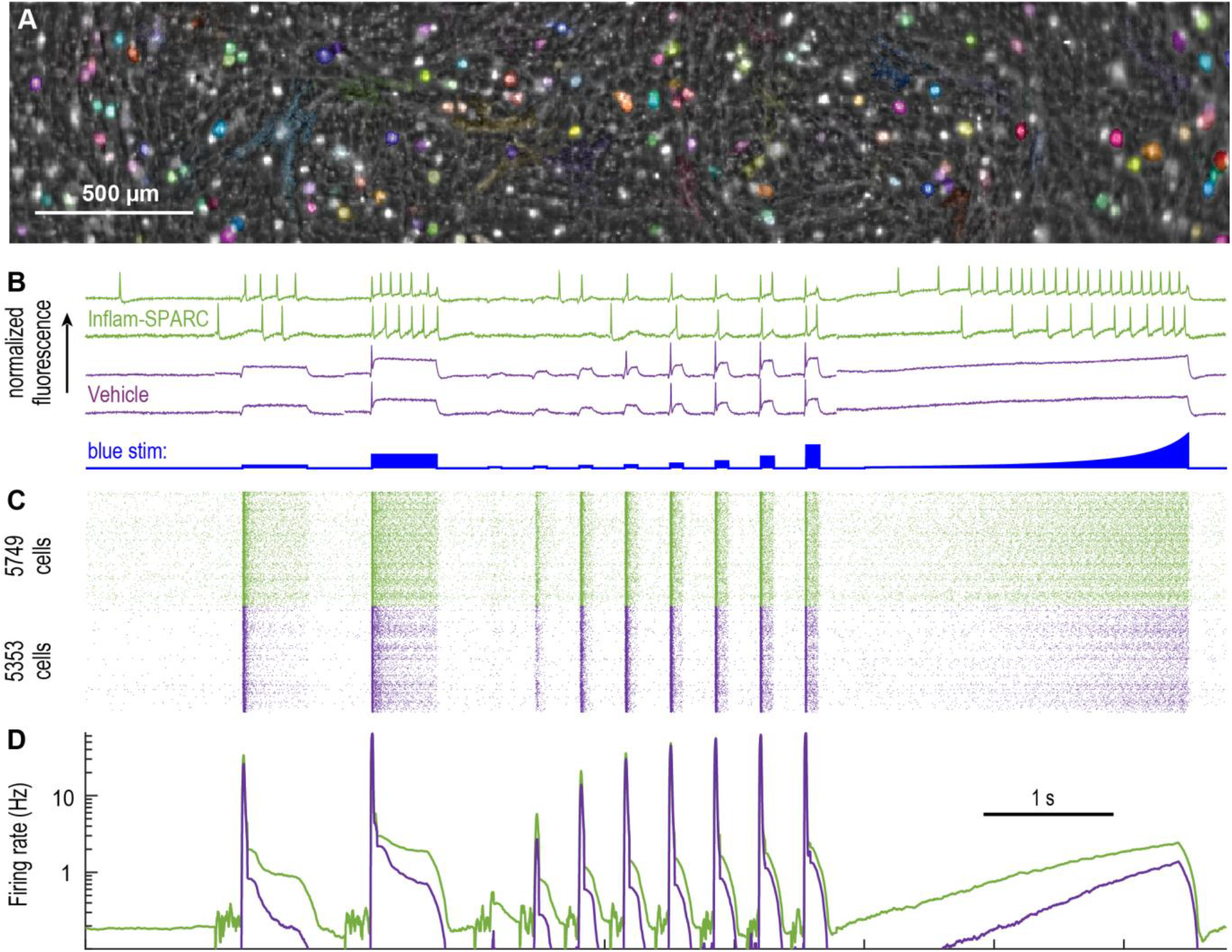
High-throughput measurements of sensory neuron excitability using all-optical electrophysiology. Rat dorsal root ganglion (DRG) neurons were dissected from P10 animals, cultured 5 days in 96-well plate format and activity recorded using the Firefly instrument. (A) Firefly image of DRG neurons with colored overlays of spiking cells identified by automated analysis. Different colors correspond to unique individual cells. (B) Example voltage recordings showing behaviors in response to step and ramp optogenetic stimuli (blue). Neurons are recorded from two conditions: vehicle control (purple) and treatment with inflammatory Sensitizing PAin Reagent Composition or SPARC (‘Inflam-SPARC’), a cocktail of inflammatory mediators designed to model inflammatory “pain-in-a-dish”. The formulation of Inflam-SPARC is (nM): 0.5 bradykinin, 100 serotonin, 100 histamine, 20,000 ATP, 10 prostaglandin E2. (C) Raster plot where each row within a color is one neuron and each point is an identified action potential. The data is pooled from 3 fields of view (FOVs) in each of 32 wells. >5,000 neurons were recorded for each condition with single cell and single action potential resolution in a single imaging session. (D) The spike rate averaged over active cells. The increased firing rate induced by the Inflam-SPARC is clearly visible, both in the raster plot and associated elevated spontaneous and evoked firing rates.

Individual cells were segmented using an activity-based algorithm. All the pixels capturing fluorescence from one neuron co-varied in time following the cell’s unique firing pattern (e.g. Fig. 1B). This covariance was used to generate a weight mask for each cell (colored masks in Fig. 1A); pixels within each mask were averaged for each frame to calculate the single-cell voltage traces. From the traces, each AP for each individual neuron in the dataset was identified and plotted in a raster graph (Fig. 1C) and firing rate was averaged over many thousands of cells (Fig. 1D).

Applying Inflam-SPARC to DRG neurons produced clear increases in AP firing during all epochs of blue light stimulation and was particularly sensitizing during sustained periods of lower-level stimulation (frequency increases from 1.09 ± 0.04 Hz in no-SPARC to 2.04 ± 0.08 Hz in Inflam-SPARC at the second long pulse with stimulus power 58 mW/cm^2^, mean ± s.e.m., p < 0.001). In addition, the optical rheobase (minimal blue light to initiate an AP) was significantly lowered by Inflam-SPARC (34 ± 2 mW/cm^2^ in no-SPARC to 9.4 ± 0.53 mW/cm^2^ in Inflam-SPARC, mean ± s.e.m., p < 0.001) indicating enhanced excitability. The example traces in Fig. 1B demonstrate the underlying variability in individual neuron behavior and illustrate the need to record from many hundreds to thousands of neurons to understand subtle pharmacological effects. These measurements were made in a 96-well plate format yielding > 300 single-cell recordings per well.

An automated analysis pipeline was developed to process the large datasets for detailed phenotype characterization. The pipeline extracted spike timing properties (e.g. spike frequency for each stimulus step, optical rheobase) (Fig. 2A) and spike shape parameters (e.g. spike width, height, after-hyperpolarization, upstroke slope, downstroke slope) (Fig. 2B) for each stimulus condition, yielding 560 parameters per neuron. We further subdivided the stimulus protocol into 46 time bins covering different portions of the protocol to capture aggregate behavior (Fig. 2C). Firing frequency and spike shape parameters were combined for each cell across each time bin, yielding 184 additional parameters that captured diverse stimulation-dependent activity patterns. The rich, multidimensional data (744 parameters in total) provided a quantitative view of neuronal behavior that can be used to assess pain disease biology and pharmacological interventions.

**Figure 2.**
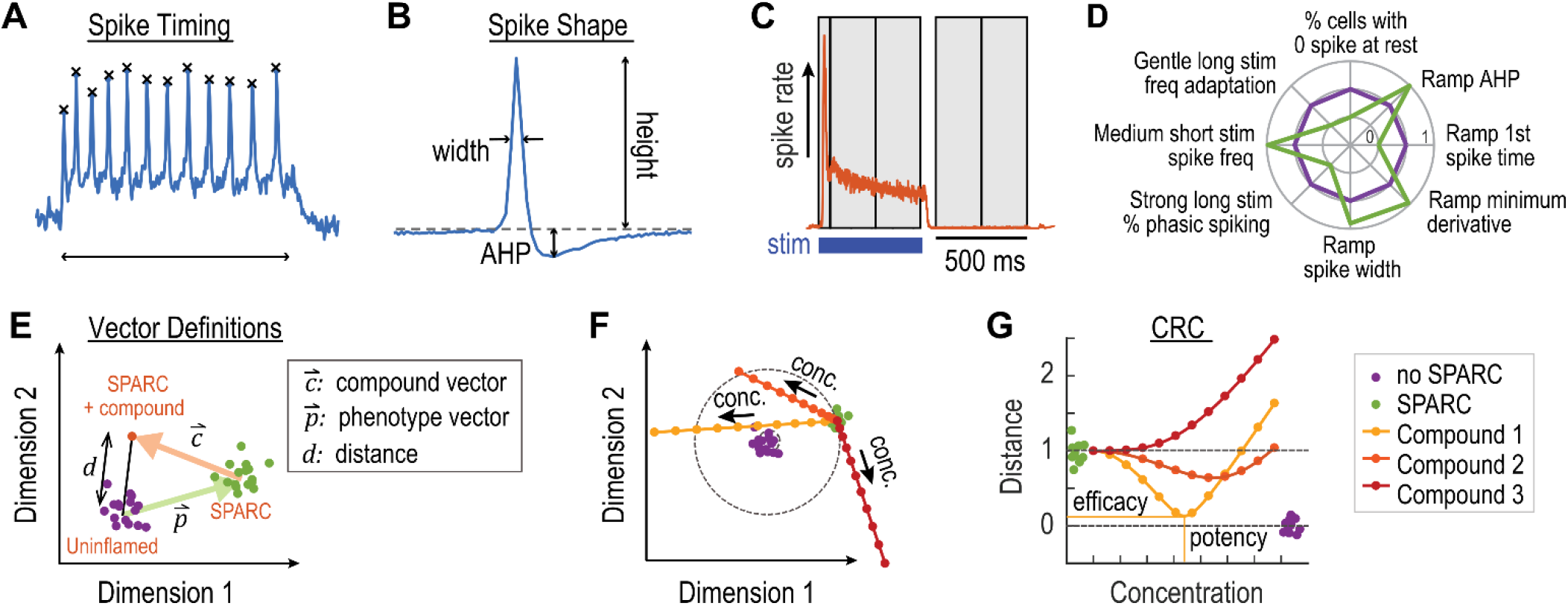
Quantitative characterization of DRG neuron activity in response to Inflam-SPARC revealed multidimensional phenotype. (A) Spike timing properties are automatically extracted including adaptation and maximum spike rate. (B) Spike shape properties are measured for each action potential, including after hyper-polarization (AHP). A total of 560 timing and shape parameters are extracted from each source trace. (C) The optogenetic stimulus protocol is broken into time bins (gray), and spike shape and timing information is aggregated from each bin, resulting in a 184-parameter vector describing neuronal function in different portions of the protocol. In total, this yields a 744-parameter dataset for each neuron. (D) The features with most significant differences after Inflam-SPARC application are identified using statistical and machine learning tools, showing a multidimensional fingerprint of neuronal behavior. Each parameter is normalized to the uninflamed condition (purple). The pain phenotype induced by Inflam-SPARC is shown in green. (E) A diagram with mock data showing examples of well location in the multidimensional phenotype space (only 2 dimensions shown for clarity). No-SPARC, uninflamed wells (purple) and SPARC-treated wells (green) form distinct clusters. A compound-treated SPARC well (orange) falls elsewhere in the space, and the Euclidean distance d between compound and no-SPARC wells relative to the length of the phenotype vector p quantifies phenotype reversal and reflects compound efficacy. The compound vector c defines the vector between SPARC-treated wells and SPARC-treated wells in response to compound treatment. (F) Concentration-response compound trajectories showing three mock examples of potential responses; yellow values move towards phenotype rescue with overshoot at high concentrations; red values move orthogonally to the ideal compound rescue vector. (G) Concentration-response curve (CRC) corresponding to mock trajectories in F. An ideal compound would reverse the inflammatory phenotype with very small distance (high efficacy) at low concentrations (high potency).

To improve data interpretability and enable robust and efficient characterization of compounds and ranking of hits from a phenotypic screen, we used statistical and machine learning tools for dimensionality reduction. We selected a set of robust, phenotype relevant parameters for each condition (see Methods: OA-SPARC Phenotyping for details), and then manually selected 8 representatives of the full phenotype for display in a “radar plot”, including 5 spike timing properties from different epochs in the blue stimulus paradigm and 3 spike shape parameters in the ramp (Fig. 2D). The difference between the Inflam-SPARC (green) and no-SPARC (purple) radar plots represents the pain phenotype. Our goal was ultimately to identify compounds that reverse the phenotype, moving the inflamed DRG spiking behavior onto that of uninflamed DRGs. Parameter values in the radar plots can provide mechanistic information on effects of SPARCs and compounds on sensory neuron function. For example, the hyperexcitability induced by Inflam-SPARC is reflected by the reduced time to the 1^st^ spike during the ramp stimulation (from 1.74 ± 0.04 s in no-SPARC to 0.83 ± 0.04 s, mean ± s.e.m., in Inflam-SPARC; Fig. 2D “Ramp 1^st^ spike time”) and the reduced percentage of silent cells at rest during periods lacking optical stimulation (from 98 ± 0.1% in no-SPARC to 91 ± 0.6%, mean ± s.e.m., in Inflam-SPARC; Fig. 2D “% cells with 1 spike at rest”).

We quantified dose-dependent effects of compounds by two numbers: a measure of phenotypic rescue (“on-target”), and a measure of orthogonal effects that did not correspond to phenotypic rescue (“off-target”, Fig. 2E). A phenotype vector, *p*, was defined as the difference between the average SPARC-treated wells and the average no-SPARC wells, in the absence of drug. A compound vector, *c*, was defined as the difference between wells treated with SPARC and drug (at a particular dose) and the average wells treated with SPARC only. The projection of the compound vector onto the phenotype vector was a measure of phenotypic rescue. We scaled this projection to create a “phenotype score” where 0 corresponded to perfect rescue, and 1 corresponded to no rescue.

To measure compound effects orthogonal to *p*, we additionally calculated the Euclidean distance, *d*, between the compound plus SPARC well (orange dot in Fig. 2E) and the average no-SPARC well (purple dots in Fig. 2E) normalized to the phenotype vector length. *d* = 0 corresponds to complete SPARC phenotype reversal with no orthogonal effects, while *d* > 1 indicates significant deviation from perfect phenotype reversal. For visualization purposes, we expressed *p* and *c* as two-dimensional vectors by projection of the 8-dimensional phenotype space into its two leading principal components (Fig. 2E, Methods). Fig. 2F shows example concentration-response trajectories. In screening applications, we prioritized phenotype reversal with near-zero “off-target” distances. An ideal compound (e.g Compound 3) would reverse the SPARC phenotype (high efficacy) at low concentrations (high potency) with no orthogonal activity (Fig. 2G).

### Establishing an *in vitro* pain model of OA using sensitized sensory neurons

We next extended the SPARC approach to model osteoarthritis pain. To this end, we developed OA-SPARC, a mixture of inflammatory mediators found in the synovial fluid of joints with chronic osteoarthritis pain [1–3,6,7,9,12,14–17,25,29,32,40,41,46,47,51,53,55–58,61,69,70,72,76,77,82– 85,87,90,91,95] (Fig. 3A, Table 1). In patients, the first step in generation of these inflammatory mediators is physical joint damage, typically from age and/or obesity[42,43]. Resultant danger-associated molecular patterns (DAMP’s), such as broken-down cartilage components and alarmins from damaged cells, are detected by local immune and repair cells. These cells in turn secrete inflammatory mediators, which accumulate in the synovial fluid, leak through the synovial membrane and modulate sensory neuron firing. OA-SPARC ingredients in the formulation meet the following criteria: (1) In humans, the mediator must be detected at increased levels in the synovial fluid of OA patient joints. (2) The associated receptor must be expressed in DRG neurons. (3) When applied to DRG neurons, the mediator must modify electrophysiology. (4) When injected in rodents, it must cause local pain. (5) After injection of the pain mediator, a pharmacological blocker of the associated receptor must alleviate pain in the injected rodent. (6) Ideally, a receptor agonist will enhance pain in the rodent. (7) Ideally, injection of the mediator will cause less pain in a mouse with receptor knockout. Table 1 provides the formulation of OA-SPARC using these criteria.

**Figure 3.**
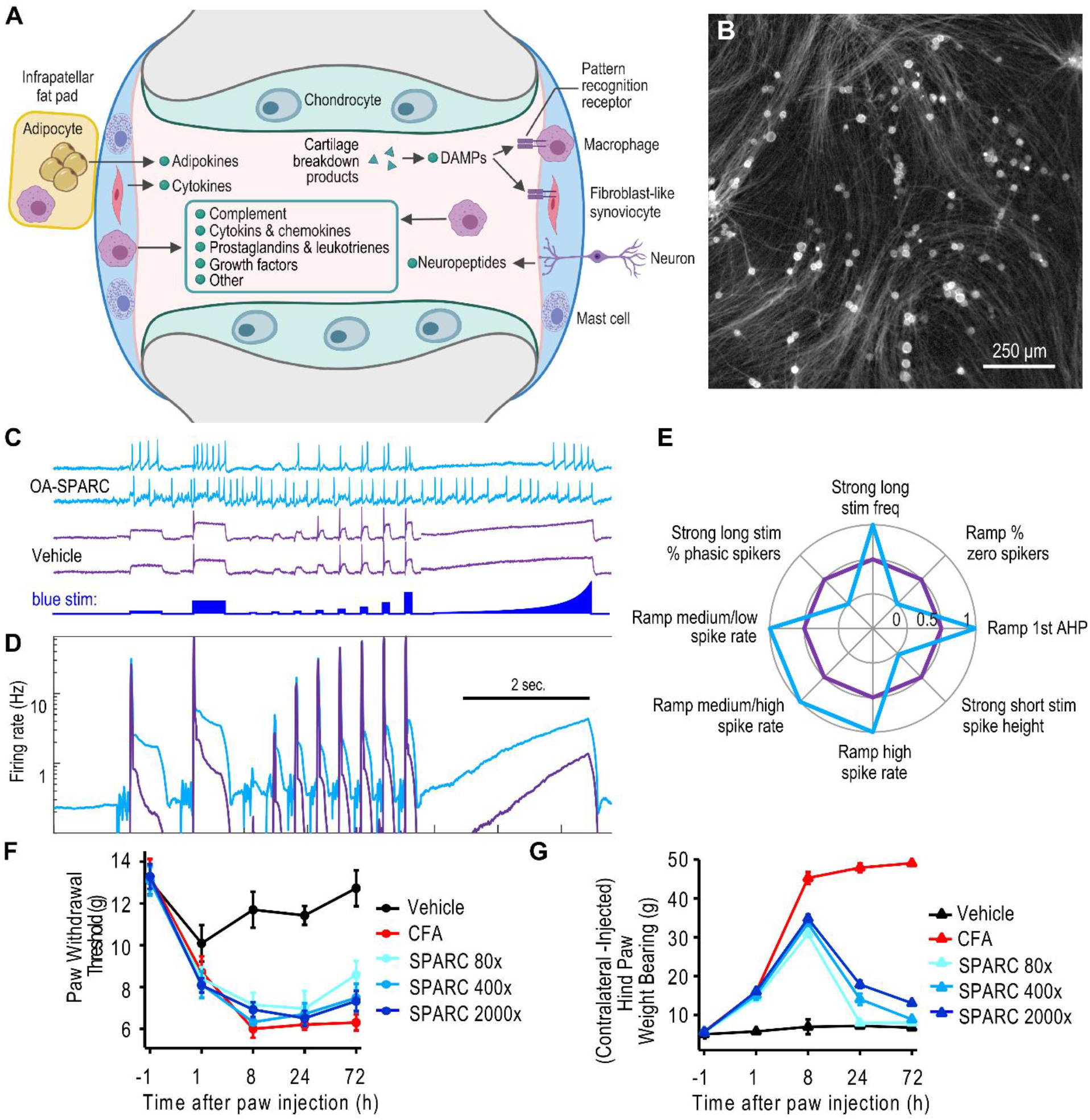
Design and validation of a SPARC for osteoarthritis (OA) pain. (A) Cartoon illustrating the complex cellular interplay in the synovial joint. In an osteoarthritic state, several inflammatory mediators are secreted into the synovial fluid of joints which leads to neuronal hyperexcitability and induced pain. (B) Rat DRGs were dissected from P10 rats, plated in 96-well plates, and transduced with viruses for optogenetic constructs 1 hr after plating. CheRiff-mOrange2 fluorescence (γ = 0.5 to highlight dim axons) is shown. The membrane-trafficked construct shows the extensive axonal processes in cultured rat DRG neurons, which contain many of the receptors that respond to OA-SPARC ingredients. (C) & (D) Neurons were treated with OA-SPARC for 24 hours. (C) Example optical physiology recordings in response to the optogenetic stimulation (blue) showed a significant increase in activity. (D) The population-averaged (>8,000 neurons in total) firing rate was increased by treatment with OA-SPARC. The SPARC induced hyperexcitability in three ways: 1. increased spontaneous firing rate, 2. higher maximal firing rate under optogenetic stimulation, and 3. firing in response to a gentler stimulus (see e.g. left edge of ramp). (E) Salient features from the 744 parameters showing a significant and stable difference between the OA-SPARC and untreated conditions were identified using statistics and machine learning, yielding a multidimensional fingerprint of neuronal behavior. Each parameter was normalized to the uninflamed condition (purple). The pain phenotype induced by OA-SPARC is shown in blue. (F) & (G) OA-SPARC was injected into the hind paws of 6-8 week old rats and leads to induced pain. Induced pain was compared against vehicle (negative control) and CFA (positive control). (F) The electronic von Frey test, where the paw was poked from below by a filament and the minimum force required to trigger paw withdrawal from the cage floor was recorded. (G) A weight bearing test, where the weight placed on both injected and un-injected hind paws was recorded. Error bars are standard deviation (SD).

**Table 1.**
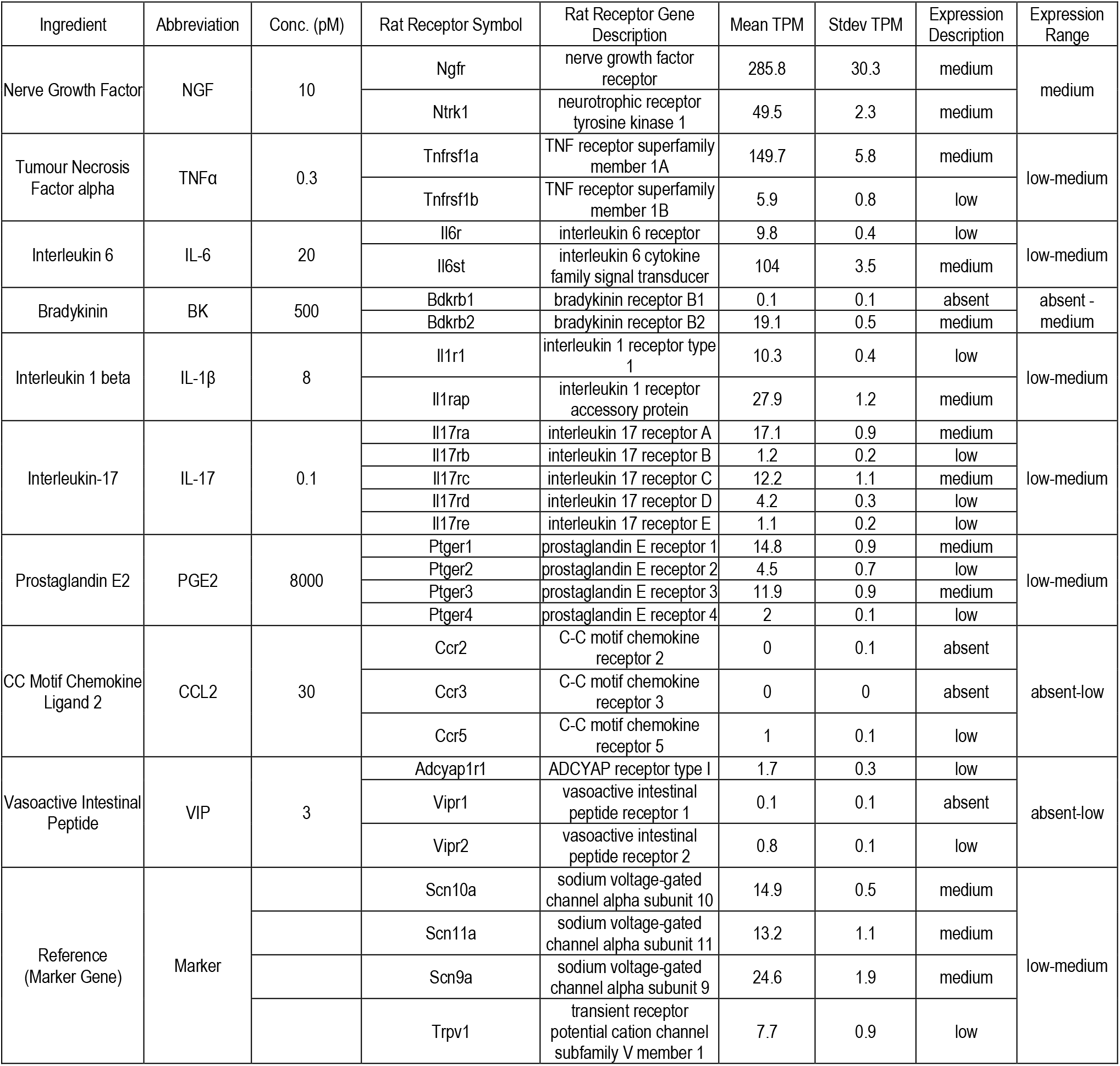
Optimized formulation of OA-SPARC. Concentrations shown are at top of the range measured in patients. Expression levels of receptor genes are quantified using transcript per million (TPM) values of these receptor genes in vehicle-treated cultured rat DRG samples. The expression of each receptor gene is categorized as absent (<0.5 TPM), low (<11 TPM), medium (<1000 TPM) or high. The expression of receptor genes for each ingredient is summarized based on the expression of each receptor gene.

Soluble factors mediate much of OA pain sensitization *in vivo*[1,25,29,33,46,53,55,56,72,83,84,91]. Approved therapeutics for OA pain have been developed targeting several components of OA-SPARC, including prostaglandin E2, Nerve Growth Factor and Tumor Necrosis Factor alpha, supporting their roles in generation of OA pain[4,5,20,23,36,74]. However, the pleiotropic mechanisms in the complex OA-SPARC mixture suggest that a blocker of just one receptor activated by the SPARC will have limited therapeutic benefit. Thus, an OA-SPARC phenotypic screen is steered toward finding compounds that modulate overall excitability.

Cultured rat DRG neurons grow extensive neuronal processes (Fig. 3B) and express many of the key receptors for OA-SPARC sensitization roles in pain signaling (Table 1). When applied to cultured DRG neurons for 24-hours, OA-SPARC induced a ~3x increase in neuronal excitability (e.g., frequency increased from 1.32 ± 0.03 Hz in no-SPARC to 3.07 ± 0.07 Hz, in OA-SPARC at the second long pulse with stimulus intensity at 58 mW/cm^2^, mean ± s.e.m., p < 0.001) (Fig. 3C, 3D). An OA-SPARC phenotype (Fig. 3E) was generated in the same way as for the Inflam-SPARC phenotype (Fig. 2D).

In addition, we validated OA-SPARC *in vivo* with measurements of pain in rats (Fig. 3F, 3G). OA-SPARC was injected into the hindpaw of rats and compared to vehicle and Complete Freund’s Adjuvant (CFA) controls. Pain was assayed by the *von Frey* test[103] (filament stiffness for first paw withdrawal) and the weight differential applied to the two hindpaws. OA-SPARC induced mechanical allodynia in paw withdrawal threshold with effect size comparable with the CFA positive control (e.g., 72-hour after injection, paw withdrawal threshold was 12.7 ± 0.9 g in Vehicle, 6.3 ± 0.4 g in CFA, 8.6 ± 0.7 g in 80x OA-SPARC, 7.5 ± 0.7 g in 400x OA-SPARC, and 7.3 ± 0.5 g in 2000x OA-SPARC, mean ± S.D.). OA-SPARC also induced statistically significant changes in weight bearing measurements at short time points (e.g., 8-hour after injection, weight bearing difference between un-injected and injected paws changed from 5.0 ± 1.1 g to 7.0 ± 1.9 g in Vehicle, 5.7 ± 0.5 g to 45.3 ± 1.5 g in CFA, 5.6 ± 0.7 g to 30.8 ± 0.5 g in 80x OA-SPARC, 5.7 ± 0.2 g to 33.6 ± 0.7 g in 400x OA-SPARC, and 5.6 ± 0.9 g to 34.8 ± 0.9 g in 2000x OA-SPARC, mean ± S.D.), indicating that OA-SPARC induced pain *in vivo*.

The final concentration for each of the OA-SPARC components was optimized with each component titrated alone and in the presence of the other components (Fig. 4). To confirm expression of the molecular targets of each of the nine OA-SPARC components along with other critical DRG genes, we performed next-generation RNA sequencing (RNA-Seq) based RNA-Seq using uninflamed rat DRG neurons. We demonstrated expression of at least one molecular target for each ligand (Table 1) in the OA-SPARC. The RNA-Seq data confirmed a medium level of expression (1000>TPM≥11 for most of the OA-SPARC receptors in comparison with a set of reference genes (*Trpv1*, *Scn9a*, *Scn10a*, *Scn11a*). Expression of receptors for CC motif chemokine ligand 2 was absent-to-low, which agreed with the small effect of this ligand when applied in isolation (Fig. 4B). The combined effect of the full SPARC was much larger than for any individual ingredient, indicating that effects were additive or possibly synergistic. Removing any individual ingredient only modestly affected the hyperexcitability phenotype, suggesting redundancy in the hyperexcitability signaling pathways (Fig. 4C). Any compounds that fully reverse the SPARC-induced phenotype will likely operate by targeting nodes where signaling paths converge (Fig. 4A). The final optimized formulation maintained clinically observed concentrations, although we applied 10x OA-SPARC to increase the signal window in screening.

**Figure 4.**
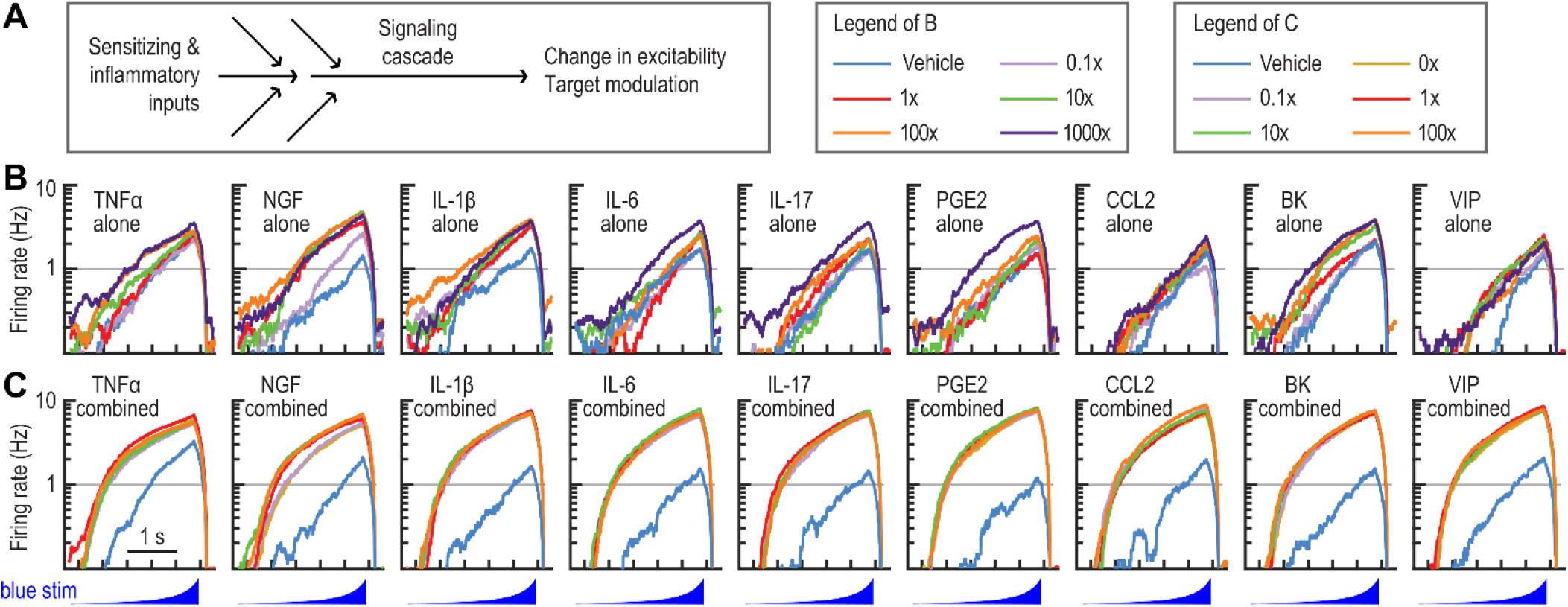
Concentration response of OA-SPARC ingredients individually and in combination demonstrated synergistic or additive effects from the combination of all components. (A) Diagram depicting inflammatory inputs from OA-SPARC that lead to a signaling cascade resulting in modulation of targets and changes in neuronal excitability. (B) The spike rate in response to a ramped stimulus (bottom). Each ingredient is titrated alone. (C) Each component is titrated in the presence of the other 8 ingredients, each at 10x concentration. The “vehicle” condition has no inflammatory mediators. All components have an effect alone, but the largest, most robust hyperexcitability phenotype is seen using the full formulation of components. Color legends for (B) and (C) are shown in the upper-right corner. 1x concentration is defined in Table 1.

### Sensory neuron OA-SPARC excitability assay validation using control pharmacology

To build confidence in the translational value of the assay, we applied pharmacological agents with defined mechanisms of action (MOA), including known analgesics, to OA-SPARC-treated sensory neurons. We started with NaV and KV7 ion channel modulators, which are known to have strong effects on neuronal excitability. Figure 5 shows the changes in spike rate properties and AP waveform as a function of concentration. A radar “phenotype” plot was generated in each case showing eight parameters identified through our analytics (see Methods: OA-SPARC Phenotyping) that displayed robust and significant effects. Plots of phenotype score and distance values versus concentration were then generated to show the extent of pharmacological rescue in each case. Phenotype and distance scores were computed using the 8 features as discussed in Methods (OA-SPARC Phenotyping). Scores were also computed using all 744 optical physiology features and the results were qualitatively similar, so we proceeded with the 8 features to simplify data analysis and visualization. Note that the phenotype score captures compound efficacy only along the OA-SPARC/vehicle control axis, while the distance score captures both on-target and orthogonal compound effects.

**Figure 5.**
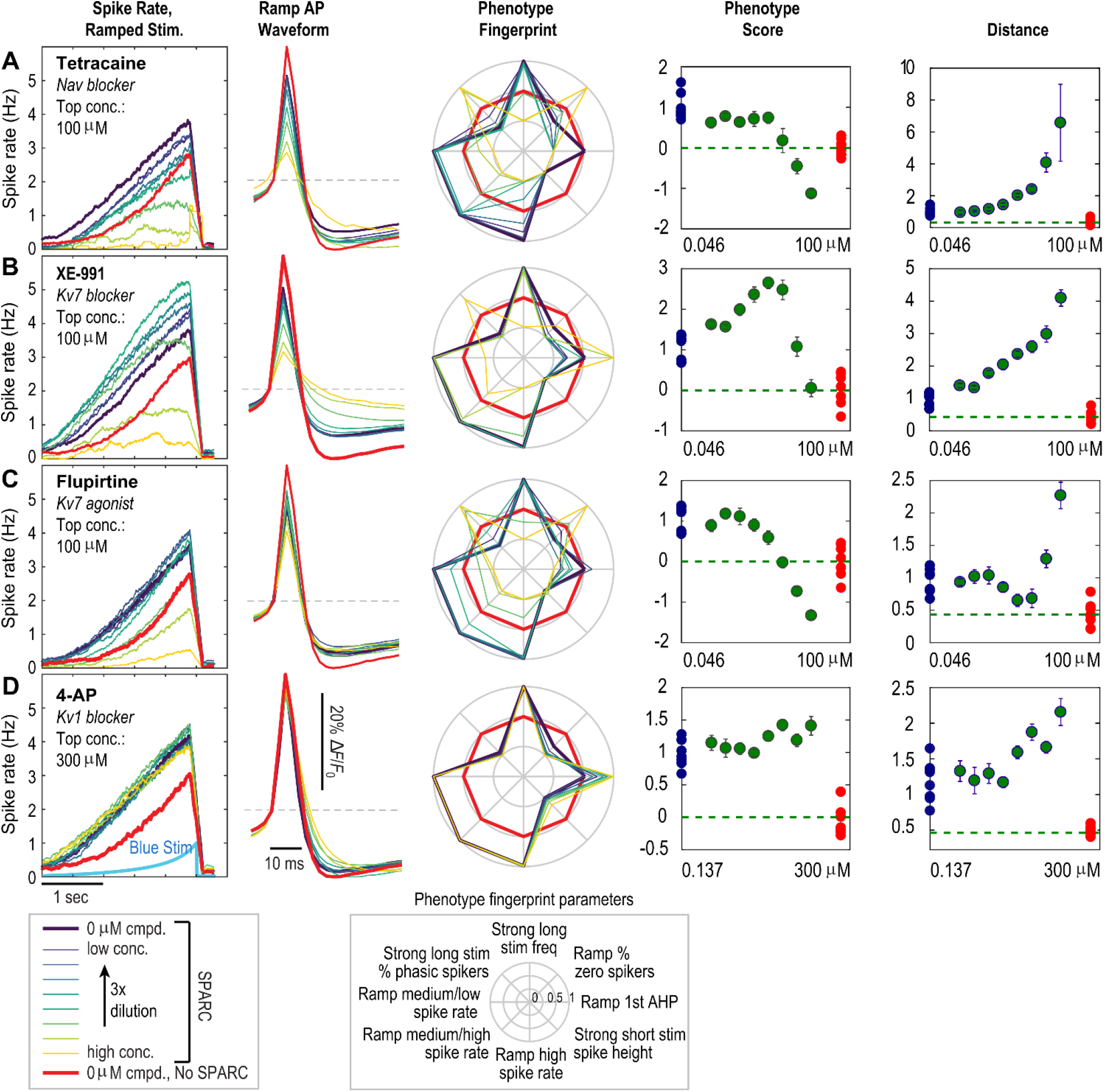
Compounds with distinct mechanisms of action showed differential phenotype reversal responses. Five plots are shown for each of the four compounds: spike rate (Hz) for ramp stimulus, AP waveform for ramp stimulus, phenotype radar plot, phenotype score and distance score. Legend for experimental condition tested and radar plot parameters are listed at the bottom. (A) Tetracaine, an approved topical analgesic, is a non-selective, activity dependent sodium channel blocker. It fully reversed the hyperexcitability phenotype at lower doses but perturbed the action potential waveform and firing at high concentrations. (B) XE-991, a Kv7 blocker, increased excitability at low doses and strongly perturbed both action potential waveform and firing rate at high doses. (C) Flupirtine, a Kv7 agonist, was formerly prescribed to treat pain in Europe. Flupirtine fully reversed the phenotype with mild effects on the action potential waveform. (D) 4-AP, a relatively nonselective Kv blocker, had mild effects on the spike rate and increased action potential width. The range of concentration, different for each compound and shown on x-axis, was selected so that published IC50 values lie in the center of the tested concentration range.

Compound effects were broadly consistent with known MOAs. Cultured DRG neurons responded robustly to sodium and potassium channel modulators. Tetracaine, a nonselective sodium channel blocker, reduced the hyperexcitability and phenotype score at low concentrations, but overcorrected the phenotype score, silenced cell firing and displayed increased distance values at high concentrations (Fig. 5A). A Kv7 inhibitor (XE-991) did not reverse the OA-SPARC phenotype (Fig. 5B), while in contrast flupirtine[27], a Kv7-activating analgesic reversed the hyperexcitability phenotype (Fig. 5C) and displayed both phenotype and distance scores indicating reversal of the OA-SPARC phenotype at lower concentrations but overcorrection of the phenotype score and higher distance values at higher concentrations, indicating possible unwanted effects at high concentrations. A nonselective potassium channel inhibitor (4-AP) did not reverse the OA-SPARC phenotype (Fig. 5D). Clinically relevant analgesics tetracaine and flupertine would be classified as active compounds in this assay while the potassium channel blockers would not be selected as antagonist hits. These results demonstrated the ability of the all-optical electrophysiology *in vitro* models to identify and characterize compounds with *in vivo* analgesic activities.

To further examine the assay performance, we evaluated 36 compounds selected based on their diverse pharmacological profiles and clinical relevance for treating chronic pain (Supplementary Fig. S1). Figure 6A shows concentration response curves for six compounds that modulate ion channel and GPCR targets. Each panel shows the distance along the phenotype recovery vector (left *y*-axis with a value of zero indicating full recovery) and the total number of APs fired per cell (right y-axis) as a general measure of neuronal excitability, as a function of compound concentration. The assay was sensitive to a variety of mechanisms including nonselective sodium channel block, Nav1.7 block, Nav1.8 block, sodium channel activation, Kv7 activation, Kv7 inhibition, KCa1.1 activation, SK1 activation, Kv1 block, KCNK3/9/18 inhibition and nociception opioid peptide (NOP) receptor agonism (Fig. S1). As expected, potassium channel agonists and sodium channel blockers typically decreased excitability, while potassium channel inhibitors and sodium channel agonists increased excitability at low concentrations. For some compounds, such as Kv7 agonist ML-213[104], both excitability and distance demonstrated compound efficacy and phenotype reversal. In contrast, Kv7 blockers increased distance and excitability measures (Fig. S1). In some cases, as for nonselective sodium channel blockers and A-803467 (Nav1.8 blocker), the distance measures decreased at low concentration, then increased at higher doses, possibly indicating exaggerated pharmacology due to on-target effects or to engagement of additional targets at higher concentrations. For other compounds, such as Cav1 agonist Bay K8644, excitability decreased at high concentrations towards the no-SPARC condition while distance increased, highlighting strong (and presumably undesired) perturbations to neuronal behavior at these concentrations. Many clinical analgesics, indicated by a *, show reversal of SPARC-induced hyperexcitability over a portion of the dosing range.

**Figure 6.**
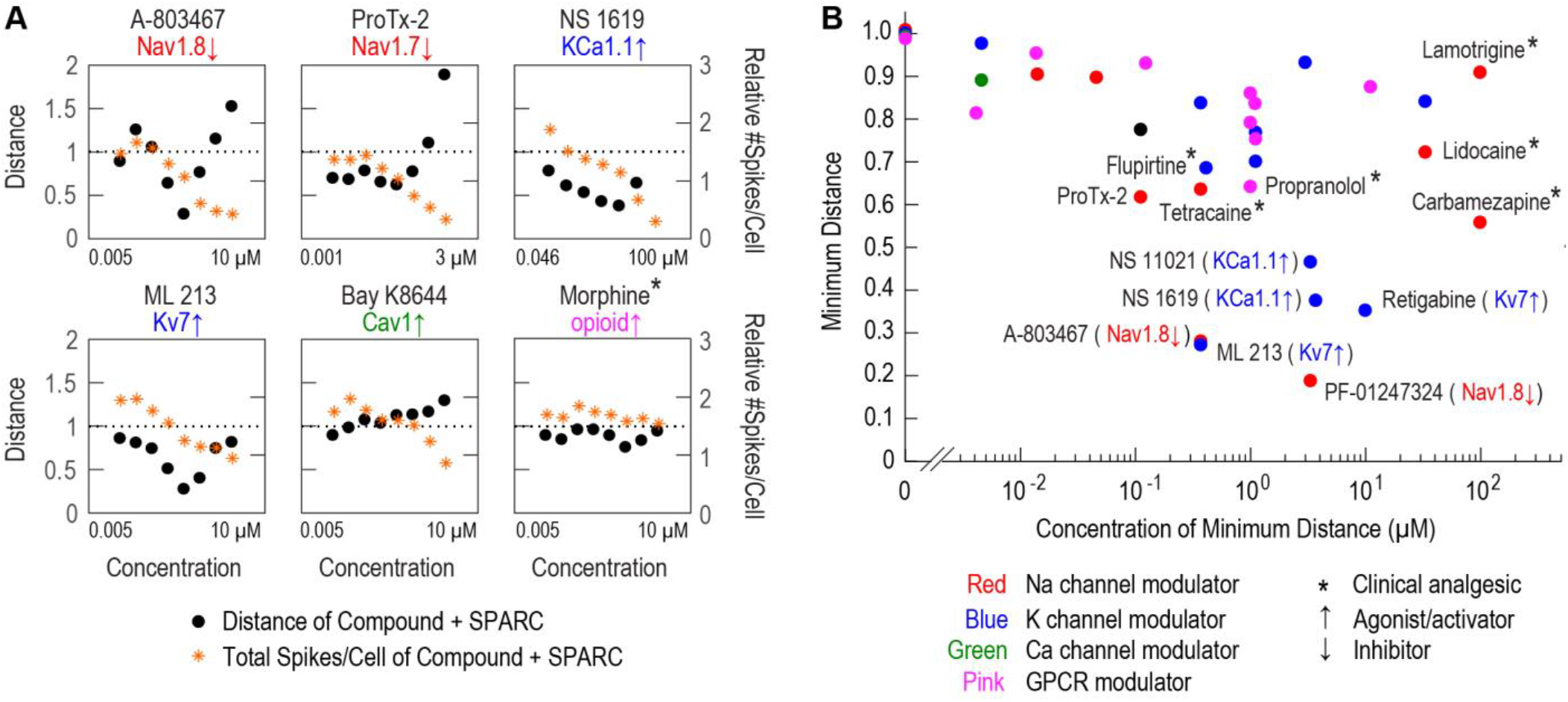
Analysis of diverse pharmacology in OA-SPARC model characterized performance characteristics of different compound mechanisms. (A) Concentration response curves for 6 compounds with diverse target mechanisms. Each panel shows an 8-point, 3x dilution series. The top concentration, different for each compound, was selected such that published IC50 values lie in the center of the tested concentration range. The y-axis on the left indicates the distance between the compound well and the average no-SPARC well as a measure of overall phenotype rescue. The y-axis on the right indicates the average number of action potentials per DRG neuron elicited by the optogenetic stimulus protocol relative to the Vehicle (no SPARC) wells. This excitability measure is reduced by many sodium channel inhibitors and potassium channel activators and increased by potassium channel inhibitors. (B) Minimum distance of 36 compounds with diverse target mechanisms are plotted versus the concentration at minimum distance.

Some clinical analgesics (Cox-2 inhibitor, opioid agonists, Cav2.2 blocker) did not affect distance or excitability measures, a consequence of low expression of the target (Fig. S1 legend) or the target’s minimal role in sensory neuron excitability. Distance measurements provided insights that were not captured by simple excitability measures, as indicated by the divergence of distance and excitability measures at high concentrations for nonselective Nav blockers. This divergence is an indicator of dose-limiting unwanted effects at high concentrations.

Therapeutic discovery efforts prioritize targets showing phenotype modulation at near-zero distances. Modulation of an ideal target would reverse the hyperexcitability phenotype with small distance (high efficacy) at low concentrations (high potency). Figure 6B plots the minimal distance of 36 compounds versus the concentration at minimal distance. Some clinically used analgesics, including carbamazepine, flupirtine and tetracaine displayed clear activity with distance values in the 0.5-0.7 range. The Nav1.7-blocking peptide ProTx-2 reversed the hyperexcitability phenotype at low concentrations, consistent with its role in pain transmission, but increased distance values at higher concentrations, possibly due to block of additional sodium channels at these concentrations. Six compounds afforded a minimum distance <0.5. Surprisingly, these six compounds fell into three MOAs: Nav1.8 inhibitors (A-803467 and PF-01247324), Kv7 agonists (ML-213 and retigabine) and KCa1.1 agonists (NS 1619 and NS 11021), suggesting Nav1.8, Kv7 and KCa1.1 channels as potential analgesic targets for OA pain, consistent with literature evidence[24,35,63,71,73,78,105]. The fact that each target was confirmed by two different compounds highlights the robust ability of the assay to identify candidate targets. Several other Nav channel inhibitors and known analgesics fell between 0.5 and 0.7 distance score.

### Optimization of OA-SPARC excitability assay for screening applications

We then sought to optimize the assay for high-throughput screening by improving the yield of active neurons and increasing the screening window. Experimental parameters that were optimized include plate type and coating protocol, OA-SPARC preparation and storage protocol, DRG dissociation protocol, plating and culture media, neurotrophic factors, cell density, viral doses, optogenetic construct promoter, imaging temperature, imaging scan pattern, imaging buffer composition and optogenetic stimulus protocol. Final conditions are described in the Methods. Rat DRG neuronal preparations were plated and treated with lentiviral particles encoding all-optical physiology components on day 0 and imaged 5-8 days later (Fig. 7A). With three dissection modules per week, we screened 1440 compounds/week. The optimized assay also had minimal time-on-microscope and plate edge effects. A full sentinel plate with zebra layout of “SPARC” and “no SPARC” wells was included in each dissection round to assess Z’ values of the OA-SPARC phenotype.

**Figure 7.**
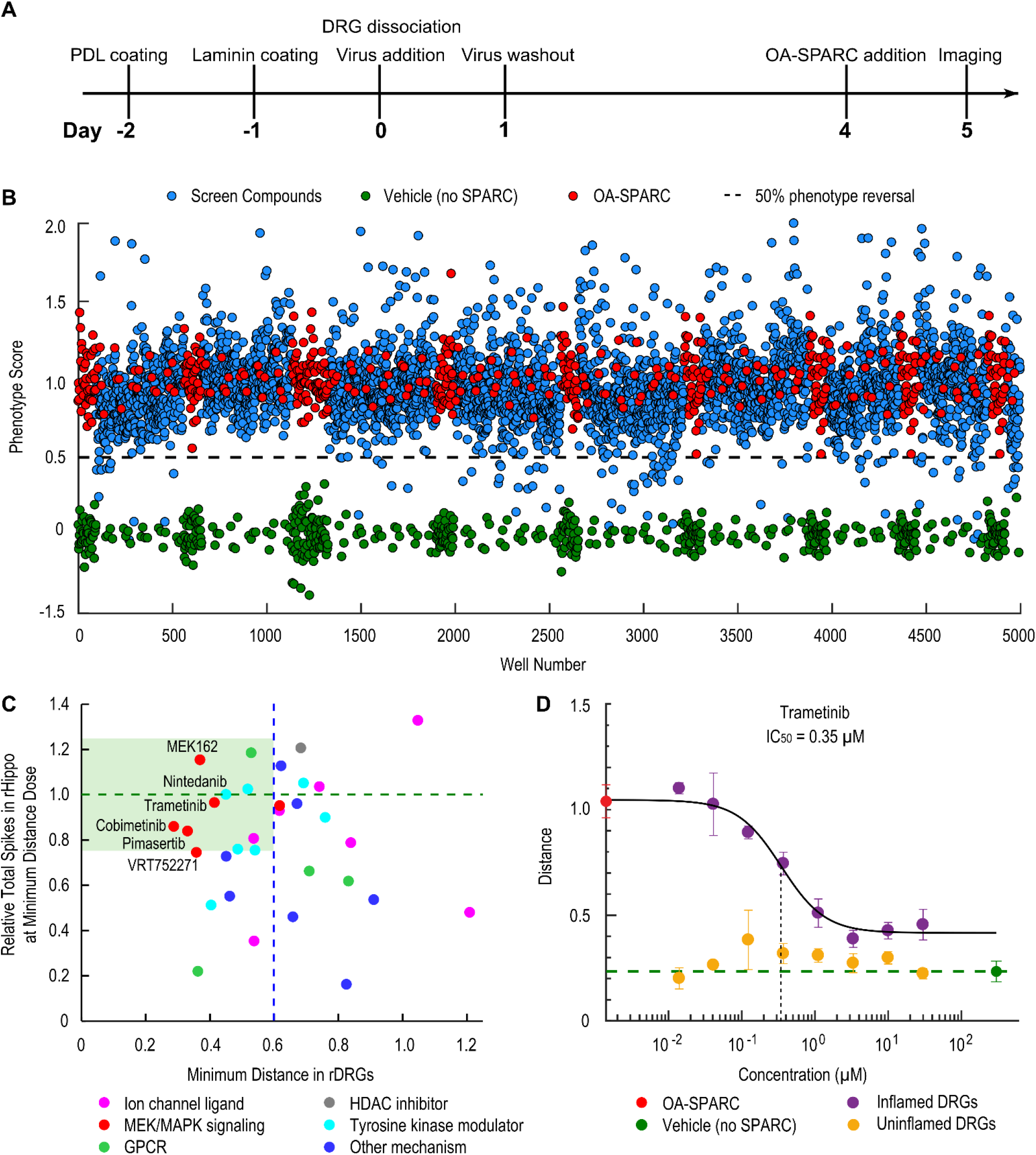
Phenotypic screen and hit validation identified known and novel mechanistic classes as potential analgesics. (A) Timeline for OA-SPARC assay. (B) Normalized phenotype score in pilot screen plates. Compounds were screened at 5μM. Hit threshold is 0.5 indicating a 50% phenotype reversal. (C) Top ranked hits were validated using an 8-point, 3x dilution series in both OA-SPARC sensitized DRG neurons and cultured rat hippocampal neurons. The x-axis indicates the minimum distance in OA-SPARC sensitized DRG neurons (efficacy). The y-axis indicates the total spikes relative to the vehicle (no SPARC) wells in rat hippocampal neurons at the minimal distance dose. Blue dashed line indicates minimum distance of 0.6, i.e. 40% phenotype reversal. Compounds within the green shaded area reversed more than 40% of the OA-SPARC phenotype with less than 25% perturbation of the spiking of hippocampal neurons, suggesting potential pain therapeutic mechanisms of interest. (D) Top ranked hits were validated using an 8-point, 3x dilutions series in both OA-SPARC sensitized DRG neurons and uninflamed DRG neurons. Shown here is a top-ranked hit, Trametinib, that reversed the phenotype in a dose-dependent manner in OA-SPARC sensitized DRG neurons but had little effect in uninflamed DRG neurons.

We evaluated DMSO tolerance to identify DMSO levels that produce < 10% changes in the assay window compared with buffer control. We found that 0.5% DMSO was tolerated for 48 hours though we proceeded with 0.1% DMSO for screening. We confirmed assay stability using three consecutive rounds of experiment. The assay achieved a stable Z’ > 0.3 which is sufficient for screening: assuming well scores follow Gaussian distributions and optimal thresholding, an assay with Z’ = 0.3 has a theoretical 77% chance of 0 false positives or negatives in a 14,000-compound screen. We also tested assay stability using IC50 values of two control compounds, tetracaine (Nav blocker) and ML-213 (Kv7 agonist). In three rounds of measurement, tetracaine and ML-213 were assayed in 8-pt concentration-response curve (CRC) experiments in quadruplicate. IC50 values of both compounds varied < 3x across three rounds (tetracaine: 2.5, 6.6, 5.3 μM; ML-213: 0.51, 0.68, 0.71 μM), confirming assay stability. The optimal screening concentration was selected by testing two screening plates randomly selected from an internal master library of ~200,000 CNS-focused, commercially available small-molecule compounds. These plates were screened at 1, 3 and 5 μM with 0.1% DMSO concentration. To rank compounds in high-throughput screening, we used the phenotype score, defined above (Fig. 2). In subsequent compound profiling in CRC experiments, we used the distance measure, *d*, to probe orthogonal effects as described in Fig. 2. We selected phenotype score = 0.5 as the hit threshold corresponding to a 50% phenotype reversal, and 5 μM as the compound concentration for screening, which maximized total number of active neurons and minimized total side effects summed over all hits.

### Phenotypic screen of annotated library

We screened approved drug and mechanistically focused libraries, including a library of ~2400 approved drugs (FDA, EMA, other agencies) assembled from four commercial libraries (Prestwick, Enzo, ApexBio and TargetMol) and a series of annotated libraries from Biomol targeting diverse neuronal proteins containing 842 additional compounds that are known to interact with neuronal signaling and regulatory mechanisms. Figure 7B shows overall assay performance with an average Z’ = 0.42 for all plates in the screen. Using the hit threshold we established during the optimization phase, we identified 133 hits including 109 antagonists that reversed the phenotype with minimal side effect scores and 24 agonists that significantly exacerbated the phenotype. The 4.4% hit rate is in line with expectations as the libraries are composed of pharmacologically active compounds.

The 133 hits from the screen were retested in duplicate at the screening concentration (5 μM) and at 1 μM. The hit confirmation rate was calculated as the fraction of compounds for which the mean of the duplicate retest measurements at 5 μM separated from control wells by at least 3 standard deviations[28]. Out of 109 antagonists selected in the screen, 45 compounds (41%) were confirmed. Out of 24 agonists selected in pilot screen, 19 compounds (79%) were confirmed. The overall hit confirmation rate was 48% (64 out of 133 compounds).

A detailed analysis of the confirmed hits revealed several common molecular pathways (e.g. sodium channel inhibitors) and candidate analgesia targets. Interestingly, six mitogen-activated protein kinase (MEK) inhibitors were confirmed as reversing the OA-SPARC phenotype, consistent with previous findings that intrathecal administration of the selective MEK inhibitor PD 198306 dose-dependently blocked static allodynia in both the streptozocin and the chronic constriction injury (CCI) models of neuropathic pain[18,101]. Trametinib, a MEK inhibitor, showed >70% reversal of the OA-SPARC phenotype with an EC50 value of ~350 nM (Fig. 7C). MEK is reported to be a key player downstream of multiple pain-related signaling pathways, including TNFα, NGF, IL-6 and PGE2[45,50,54,93,94,100]. Our identification of MEK as a target supports our hypothesis that compounds acting on convergent points of multiple pathways are more likely to demonstrate greater phenotype reversal.

Two histone deacetylase inhibitors (HDACis) were also confirmed, consistent with their interference role in the epigenetic process of histone acetylation, and analgesic properties in models of chronic inflammatory pain[21,102]. Nine compounds were confirmed to exacerbate the OA-SPARC phenotype (Fig. S2A). These compounds may provide insight into potential pathways involved in generating a proalgesic condition, and as a signal of potential off target effects of known drugs. For example, chemotherapeutics cisplatin and carboplatin are well known to induce peripheral neuropathy in some patients[13,86].

The 33 top-ranked antagonist hits (Supplementary table 1) that were commercially available were reordered as fresh samples from alternative vendors when possible and were then counter-screened in two additional optical electrophysiology assays to identify potential off-target effects in the central nervous system (CNS) and in un-sensitized DRG neurons. To detect undesired activity in the CNS, we counter-screened the 33 hits in 8-point CRC in cultured rat hippocampal neurons, using an optical excitability assay (Methods). Figure 7C compares the efficacy in the OA-SPARC DRG assay (x-axis) versus CNS effects (y-axis). Ideal compounds reside in the shaded region and show clear OA-SPARC phenotype reversal (low minimal distance values) with minimal effects on AP frequency in hippocampal neurons. CRC results provide a means to rank hits, to evaluate assay reliability and hit criteria and to validate whether hits impacted excitability via an OA-SPARC related mechanism.

We then counter-screened the 15 antagonists which showed minimal CNS perturbation in un-inflamed DRG neurons, to assess specific reversal of the OA-SPARC phenotype and to search for compounds with minimal effects on normal sensory activity. Trametinib, a MEK inhibitor, showed no effect on the parameters used to define the OA-SPARC phenotype in DRG sensory neurons without OA-SPARC sensitization (Fig. 7D), indicating a selective effect on OA-SPARC induced hyperexcitability.

Nineteen agonists that enhanced the OA-SPARC phenotype were evaluated in CRC to investigate potential mechanisms associated with pain signaling pathways and to identify potential adverse effects of approved drugs. Figure S2 presents a list of the nine top ranked agonist hits and CRC data for four agonists tested in the absence and presence of OA-SPARC. All four compounds were active in both the absence and presence of OA-SPARC. The concentrations for half maximal effects in the DRG assay for some compounds were within clinically relevant ranges of *in vivo* exposure. In some cases, the agonist effects on DRG activity may not be coupled to the primary pharmacological mechanism; loratadine is a widely used antihistamine (H1 receptor antagonist), which also blocks KCNK18, a potassium channel involved in pain[10]. Further exploration of these hits may yield new information on pathways involved in generating or reducing hypersensitivity in sensory neurons for OA pain.

## Discussion

Over recent decades, the pharmaceutical industry has focused predominantly on target-based approaches to develop new pain therapeutics because associated high-throughput methods enable efficient screening of large libraries and rapid characterization of analogues[11,99]. Additionally, knowing the target aids rational molecular design during medicinal chemistry optimization, helps guide preclinical safety studies and enables development of target engagement strategies in clinical trials. Despite these advantages, the scarcity of new classes of analgesic drugs argues for establishing alternative development pipelines, including phenotypic screening[39].

Phenotypic assays historically played an important role in pain drug discovery. The widely used pain therapeutics gabapentin (Neurotin) and pregabalin (Lyrica) were initially discovered as anticonvulsant agents using phenotypic screens prior to identification of the α_2_δ-_1_ subunit of voltage-gated calcium channels as the molecular target[26]. Proven success of phenotypic approaches is leading to a resurgence in use for some therapeutic areas[22,92,107]. Because compounds are screened in a disease relevant cellular system, measured responses can be more predictive of clinical results. Compounds that strongly modulate sensory neuronal function *in vitro* are likely to modulate function of the same neurons *in vivo*. For example, cultured neurons can contain appropriate protein splice isoforms, expression levels, post-translational modifications and interacting partners, which can dramatically affect drug performance.

Phenotypic screening also casts a broad net: there are over 12,000 mRNA transcripts expressed in the human DRG, over 3,000 of which are elevated relative to fibroblasts or Schwann cells[75]. 141 protein-coding genes are expressed at elevated levels relative to central nervous system (CNS) tissue[75] and so are potential drug targets with lower risk for on-target toxicity. A phenotypic screen will be sensitive to modulation of many of the targets. Another key advantage is that phenotypic assays can detect pleiotropic drugs[59], which have the potential for greater efficacy than a single-target compound due to synergistic effects. By using a primary phenotypic screen in DRG neurons and phenotypic counter screens in CNS neurons and cardiomyocytes, the approach will also be sensitive to safety issues arising from pleiotropic effects. An example pleiotropic drug is clozapine, a highly efficacious atypical antipsychotic that interacts with multiple receptors[68].

The probability of success for a phenotypic screen depends on the translatability of the assay to clinical outcomes, both the cellular model and the assay readout. Our cellular “*pain-in-a-dish*” model for OA-pain induced rat DRG neuronal sensitization uses OA-SPARC, a novel formulation of inflammatory mediators present at elevated levels in arthritic joints. OA-SPARC application drove a strong *in vitro* hyperexcitability phenotype in the cultured DRG neurons and caused pain *in vivo* when injected into a rat paw. The use of a rodent DRG model is supported by the fact that 128 of the 141 protein-coding genes that are expressed at elevated levels relative to central nervous system (CNS) in humans have conserved orthologs in mice[75]. Human induced pluripotent stem (IPS) cell derived sensory neurons are emerging as tools for pain disorder modeling induced by genetic mutations[60,65] but so far have limited applications in modeling pain induced by inflammatory mediators, including OA. To build confidence in the primary rat neuronal model, we pharmacologically validated the model by showing hyperexcitability reversal with clinical analgesics and sensible responses to a battery of potent and selective compounds with diverse targets.

The OA-SPARC model has been enabled for drug discovery by pairing with a high-throughput all-optical electrophysiology platform[98]. Manual patch clamp electrophysiology provides exquisite sensitivity but at the expense of throughput. Automated patch clamp studies provide detailed compound mechanisms, but is limited by high cost, modest throughput and does not work well in intact cultured neurons. Calcium imaging provides high throughput by leveraging optical methods but lacks targeted stimulation and temporal resolution required to detect action potentials. Similarly, fluorescence plate readers provide high throughput, but are associated with low temporal and spatial resolution and thus provide limited mechanistic information. The resulting translational challenge for pain drug discovery and for CNS-based disorders more broadly is significant, necessitating new approaches to bridge this important gap.

The combination of enabling screening technology and pain-relevant *in vitro* models provides promising opportunities for identification of improved therapeutics and exploration of pain signaling pathways. Because this platform is compatible with primary DRG neurons from different species, it is feasible to validate the key discoveries made in rodents using non-human primate and human primary DRG neurons. Recent progress in development of iPS cell-derived sensory neurons[80] can provide an avenue to large scale generation of relevant human cell types to enable high throughput phenotypic screening for pain therapeutics to additionally provide a scalable means to evaluate and rank compounds for sensory cell activity as part of target-based pain therapeutic development projects. Lastly, we anticipate that other forms of chronic pain induced by defined mixtures of inflammatory mediators, such as those secreted by tumors in cancer pain, may be modeled by taking a similar approach as the one described in this study.

## Methods

### Solutions

#### Dissection media

Calcium and Magnesium Free HBSS (Fisher Scientific) supplemented with 10 mM HEPES (Fisher Scientific) and 0.5x Penicillin-Streptomycin Solution (Corning), pH 7.35.

#### Plating media

DMEM/F12 medium (Fisher Scientific) supplemented with 10% heat-inactivated FBS (VWR), 10 mM HEPES (Fisher Scientific), and 0.5x Penicillin-Streptomycin Solution (Corning), pH 7.35.

#### Culture media

BrainPhys Neuronal media (STEMCELL Technologies) supplemented with 1x N2 Supplement-A (Fisher Scientific), 1x SM1 Neuronal supplement (Fisher Scientific), 1x GlutaMAX (Fisher Scientific), 20 nM Trans-Retinel (Sigma-Aldrich), 0.5x Penicillin-Streptomycin Solution (Corning), 2 ng/mL mouse GDNF (Sigma-Aldrich), 4 ng/mL human NT-3 (Sigma-Aldrich), and 1 μg/mL laminin (Fisher Scientific), pH 7.35.

#### Imaging buffer (in mM)

115 NaCl, 18 Na gluconate, 3 KCl, 1.1 CaCl_2_, 1 MgSO_4_, 0.5 Na_2_HPO_4_, 0.45 NaH_2_PO_4_, 4 D-Glucose, 10 HEPES, 0.15 Na pyruvate, pH 7.35.

### Preparation of cultured DRG neurons

#### Plate preparation

Greiner cycloolefin half-area 96-well plates (Greiner Bio-one, Item No.: 675801) were plasma-treated for 3 minutes at 0.26 Torr with air (oxygen) plasma to make the surface hydrophilic. Plates were then sterilized by exposure to UV light in a tissue-culture hood for 30 minutes. Two days prior to plating of DRG neurons, wells were coated with 100 μg/mL poly-d-lysine (Sigma-Aldrich) in water and incubated at 4°C overnight. One day prior to plating of DRG neurons, wells were washed with PBS and coated with 20 μg/mL laminin (Fisher Scientific) in PBS and incubated at 4°C overnight. On the day of plating of DRG neurons, plates were retrieved and equilibrated to room temperature. Immediately prior to plating DRGs, laminin was aspirated, and wells were washed with plating media. From one rat litter, we routinely plated seven full 96-well plates.

#### Cell preparation

DRG neurons were prepared from Sprague Dawley rats, postnatal day 10. Animals were anesthetized by using CO_2_ inhalation and decapitated, and ganglia were removed. Dorsal root ganglia from one litter of 10 pups were treated at 37°C for 30 min in 2 mg/mL collagenase (Type IA; Sigma-Aldrich) and 5 mg/mL dispase II (Sigma-Aldrich) in dissection media and then washed with and placed in plating media. Individual cells were dispersed by trituration with a fire-polished Pasteur pipette and filtered through 100 μm cell strainer. Cells were then counted and plated at a density of 2,000 cells/well on Greiner half-area 96-well plates with a cyclic olefin copolymer (COC) bottom (Greiner Bio-one, Item No.: 675801) in plating media. COC, which was optimized for ultraviolet light applications, also makes substrate autofluorescence and laser-generated sample heating negligible upon red laser illumination[67]. The small well area conserves cells (2k/well) and enables imaging of nearly the full well area. During cell preparation and culture maintenance, we used semi-automated culture with the Integra VIAFLO 96-channel pipettor, which reproducibly defines pipette location (*x*, *y*, *z*) and injection/aspiration velocity[67].

#### All optical electrophysiology construct transduction

One hour after cell plating, Lenti-X Concentrator concentrated lentiviruses including CaMKII-CheRiff-EBFP2 and CaMKII-QuasAr3-Citrine were added for transduction. Cells were then incubated in a cell culture incubator with 5% CO_2_ at 37°C overnight. 16-18 hours after transduction, virus was removed and cells were fed with culture media until compound treatment.

#### Cell culture maintenance and compound treatment

OA-SPARC stock solution was prepared at 1000x (see recipe in Table 1), aliquoted and stored at −80°C. Compounds were prepared at 200x in DMSO and stored at −20°C. 24-hour prior to imaging, OA-SPARC and compounds were diluted in culture media and added to the cells at the assay concentration. 30 minutes before imaging, culture media was aspirated from the wells and OA-SPARC and compounds were prepared in imaging buffer and added to the cells using Perkin Elmer JANUS Mini automated liquid handling system.

### All optical electrophysiology imaging

Imaging experiments were performed 5 days after DRG neuron plating into 96-well plates. Prior to imaging, the cells were first washed with imaging buffer then treated with OA-SPARC and tested compounds using a PerkinElmer Janus Mini liquid handling robot. Cells were then incubated for 30 min at 27°C. Imaging experiments were performed on the Firefly microscope equipped with an automated stage (Ludl BioPrecision2, Ludl Electronic Products, Hawthorne, NY) at 27°C. Briefly, QuasAr3 voltage sensor were illuminated by two red lasers (638 nm, 8 W, DILAS-Coherent, Santa Clara, CA) was homogenized and refracted onto the sample via near-TIR illumination (NA 0.4, UPlanSApo 10×/0.4; Olympus, Shinjuku City, Tokyo) at light intensity around 120 W/cm^2^. Blue light for optogenetic stimulation was projected by a high-power LED (462 nm, 38W, Luminus Devices Inc., Sunnyvale, CA) via a fast digital mirror device (DMD) (Vialux, Chemnitz, Germany) through an objective lens (NA 0.5, 2x/0.5 MV Plapo 2XC, Olympus, Shinjuku City, Tokyo) onto the sample. QuasAr3 fluorescence was filtered by a 736/128 nm bandpass filter and detected by a CMOS camera (ORCA-Flash4.0; Hamamatsu, Bridgewater, NJ). To achieve a frame rate of 500 Hz, data were acquired with a center camera chip of 800 × 160 pixels at 2 × 2 pixel binning, which corresponds to the 4 × 0.8 mm recording area in the well. Scanning progressed from well A1 to well A12, then B12 to B1, then C1 to C12, and so on until in well H1, in a column-wise serpentine pattern, then reversed from H1 to H12, then G12 to G1, then F1 to F12, and so on until back to well A1, in a reversed column-wise serpentine pattern, then in the column-wise serpentine pattern again, i.e. from A1 to A12, then B12 to B1, then C1 to C12, and so on until in well H1. Using this serpentine-reversed serpentine-serpentine pattern reduced the time-on-microscope effect. Each well was typically stimulated with a protocol consisting of two long pulses (11.25 mW/cm^2^, 58 mW/cm^2^, 500 ms each), eight short pulses (2.44 mW/cm^2^, 5.5 mW/cm^2^, 9.43 mW/cm^2^, 14.67 mW/cm^2^, 22 mW/cm^2^, 33 mW/cm^2^, 51.33 mW/cm^2^, 88 mW/cm^2^, 100 ms each), followed by a linear conductivity ramp pulse (0.125 mW/cm^2^ to 30 mW/cm^2^ in 2.5 s) of blue light (see Fig. 1B). A complete 96-well plate required 67 min to scan using this protocol.

### Small molecule library

A library of approximately 2400 approved drugs was assembled from four commercial sources (Prestwick, Enzo, ApexBio, and TargetMol). These drugs had been approved by regulatory agencies in the US, Canada, European Union, Japan and other countries. Due to duplications across individual libraries, this collection of approved drugs was contained in ~3400 individual wells. This library was supplemented with a series of annotated libraries from Biomol containing 842 additional compounds that target a variety of CNS signaling/regulatory mechanisms and expressed proteins including endocannabinoids, ion channel ligands, adrenergic ligands, dopaminergic ligands, opioid ligands, cholinergic ligands, histaminergic ligands, ionotropic glutamatergic ligands, metabotropic glutamatergic ligands, GABAergic ligands, purinergic ligands, kinase inhibitors and phosphatase inhibitors.

### OA-SPARC *in vivo* validation

This study was conducted at Pharmaron Beijing Co., Ltd. (China). SD rats with age 6-8 weeks were kept in laminar flow rooms at constant temperature and humidity with 3 animals in each cage. Animals were housed in a polycarbonate cage and in an environmentally monitored, well-ventilated room maintained at a temperature of (22±3°C) and a relative humidity of 40%-80%. Vehicle (PBS), Complete Freund’s Adjuvant (CFA) or OA-SPARC dissolved in PBS was injected into the left hind paw. Von Frey threshold and Weight Bearing measurements were performed five times during the study including baseline prior to dosing, and 1 hour, 8 hours, 24 hours and 72 hours post dosing to evaluate the effects of each test agents. Mechanical allodynia of the left hind paw was measured during the course of study by determining withdrawal thresholds to an electronic Von Frey filament (Bioseb, France). Rats distributed body weight unequally on Dose-Infection and contra-lateral paws, which were measured by weight balance changing instrument (YLS-11A, JinanYiYan Science and technology development co., LTD). The animals were tested to register the weight load exerted by the hind paws by means of a force plate inserted in the floor.

### Analysis

#### Analysis pipeline

Movies were recorded serially from each plate and the typically 100s of Gigabytes of data were transferred to Amazon Web Services (AWS) for storage and parallelized analysis. Using a watershed algorithm based on the movie mean frame, the field of view (FOV) is divided into regions which are individually segmented to find sources. Patch-wise Principal Component Analysis (PCA)/Independent Component Analysis (ICA)[62] was employed to find spatiotemporally correlated fluorescence signals. The patches were then merged and sources were combined or split. The temporal covariance of pixels for each cell was used to generate a weighted mask; pixels were averaged for each frame in the movie to calculate the single cell voltage-time traces. Each trace was corrected for photobleaching and crosstalk, and spike detection was performed to identify action potentials in the traces. Each action potential was parameterized and dozens of spike shape parameters are extracted, furthermore, spike frequency and timing properties are extracted, yielding 560 parameters per cell. Cells were scored based on morphological properties such as size and connectivity of the cell bodies as well as signal-to-noise ratio of the detected spikes, and cells with a low score were discarded. Results were downloaded into a database, from which feature tables are generated for further analysis.

#### Time bin analysis

Based on the diverse stimulation protocol, each movie was split into 46 time bins covering different portions of the protocol and firing frequency as well as three spike shape parameters (spike height, width and after-hyperpolarization) were aggregated for each cell across each bin. This yielded 184 additional features that captured diverse aspects of cellular behavior under various stimulation scenarios.

#### Inflam-SPARC exploratory analysis

32 wells each of Inflam-SPARC and vehicle conditions were available for each sentinel plate. A t-test was performed for each of the 744 features (560 features from analysis pipeline and 184 features from time bin analysis) and the features were ranked based on most significant difference between the vehicle and Inflam-SPARC (Inflam-SPARC) groups. A significant and diverse subset of 8 features was manually selected for visualization of the effect of Inflam-SPARC on the cell population.

#### OA-SPARC Phenotyping

A multi-step statistics and machine learning approach was used to define a disease model for osteoarthritic pain consisting of 8 parameters. 5 rounds of 3 plates with 32 wells each for diseased and healthy conditions were used in the phenotyping process. First, for each of the 744 features, linear mixed effects models were used to estimate p-values for the difference between diseased and healthy conditions in a way that was sensitive to hierarchical sources of experimental variance. A false discovery rate correction was performed, and the significant features were passed on into a LASSO regression, which automatically performed feature selection by setting coefficients of non-relevant features to zero. Around 40 features were obtained, and a further manual selection for lowest correlation and good interpretability while keeping a stable phenotypic window yielded 8 features to describe the OA-SPARC phenotype. These features were combined using a principal components analysis to account for covariance; then we used vector decomposition to compute a single disease score: in the principal component space, an *average disease vector* was defined that points from the average of all OA-SPARC wells to the average of vehicle (no SPARC) wells. The position vector for each individual well was projected onto the average disease vector, and the length of the projection was called the on-target score, describing activity along the desired direction in the principal component space. The length of the portion orthogonal to the average disease vector was called the off-target orthogonal score. A total distance score was also computed, defined as the distance of a given well from the average of vehicle (no SPARC) wells, and is effectively Mahalanobis distance. The Z’ factor for the screen was defined as Z’ = 1– 3(σ_p_+σ_n_)/1 (μ_p_-μ_n_) |, where σ_p_ is the standard deviation of the vehicle (no SPARC) wells, σ_n_ is the standard deviation of the OA-SPARC sensitized wells. μ_p_ is the mean of the vehicle (no SPARC) wells and μn is the mean of the OA-SPARC sensitized wells. The on-target score consistently exhibits a Z’ greater than 0.2 for the phenotyping plates and is used as the primary metric in the high-throughput drug screen and hit confirmation. The total distance score was used to evaluate confirmed hits in dose response experiments.

## Acknowledgements

We thank Ted Brookings and Trinh Nguyen for early contributions to assay establishment and analysis tools. This work was funded in part by NIH grant 5R44AR074820. PL, HZ, CW, MP, SR, CL, JJ, JG, JF, GL, DZ, NB, AB, RC, AE, LW, GD and OM are current or former employees of Q-State Biosciences and may hold stock options in Q-State Biosciences. AC is a co-founder of Q-State Biosciences and may hold stock options in Q-State Biosciences.

