## Supplemental Table 1 for "A phenotypic screening platform for chronic pain therapeutics using all-optical electrophysiology"

Supplementary figure 1

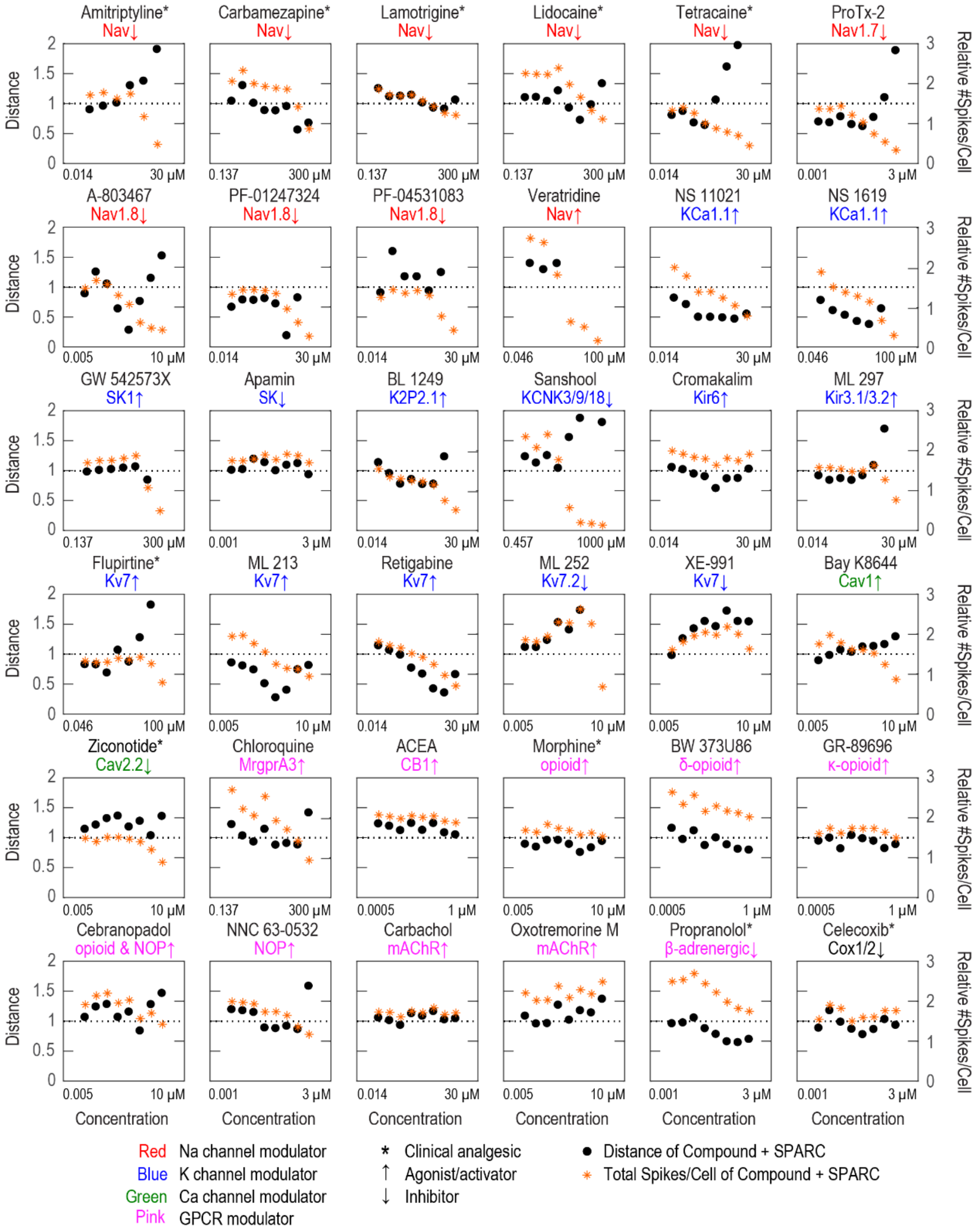


**Supplementary figure 1 | Examples of diverse pharmacology using OA-SPARC phenotypic assay.** Concentration response curves for 36 compounds with diverse targets. Each panel shows an 8-point, 3x dilutions series. The top concentration, different for each compound and shown on the right end of the x-axis, was selected so published IC50 values lie in the center of the tested concentration range. The y-axis on the left indicates the distance between the compound well and the average Vehicle (no SPARC) well. The y-axis on the right indicates the average number of action potentials per DRG neuron elicited by the optogenetic stimulus protocol relative to the Vehicle (no SPARC) wells. This excitability measure was reduced by many sodium channel inhibitors and potassium channel activators and increased by potassium channel inhibitors.

Supplementary figure 2

**A**

| Enhancers | MOA | Maximum phenotype score | Conc. of half-maximum effect (μM) | Clinical/in vivo dose (μM) |
|---|---|---|---|---|
| GSK2126458 | PI3K/mTOR inhibitor | 2.71 | 0.23 | 0.15 |
| Daclatasvir | HCV NS5A inhibitor | 1.45 | 2.50 | 0.25 |
| Mitotane | Steroidogenesis inhibitor | 2.34 | 1.95 | 43.74 |
| Beclomethasone dipropionate | Glucocorticoid receptor agonist | 1.46 | 1.87 | 0.00 |
| Vilazodone hydrochloride | 5-HT receptor inhibitor | 2.52 | 3.65 | 0.33 |
| Conivaptan HCl | Vasopressin receptor antagonist | 2.40 | 5.33 | 1.16 |
| Rimonabant hydrochloride | Cannabinoid receptor antagonist | 1.61 | 0.37 | 0.39 |
| Loratadine | H1 histamine antagonist, KCNK18 inhibitor | 1.82 | 0.65 | 0.01 |
| Cisplatin | Inhibits DNA synthesis | 1.25 | 2.28 | 35.99 |

**B**

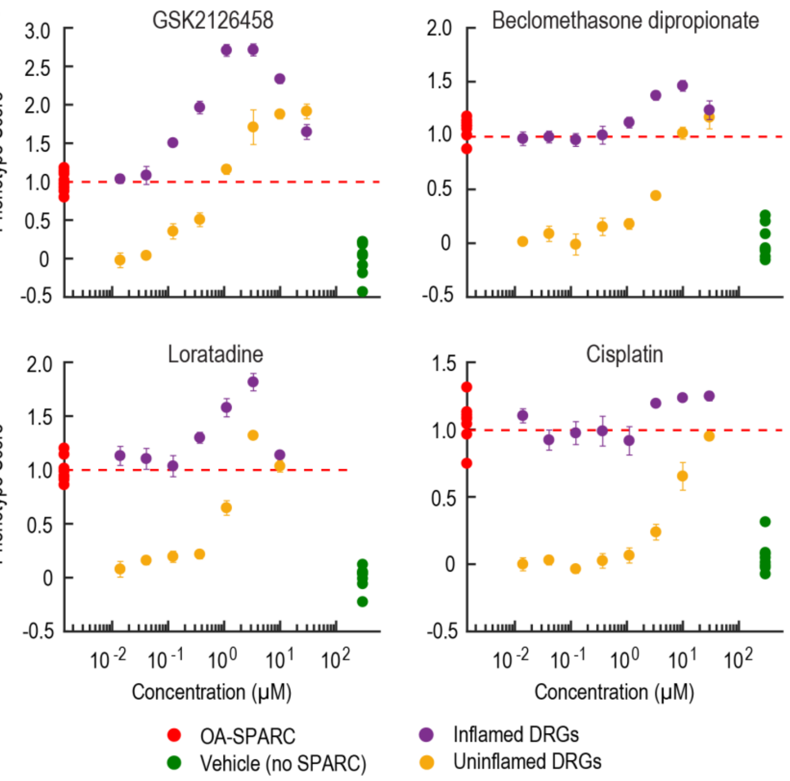

**Supplementary figure 2 | Hits identified that exacerbate OA-SPARC phenotype include known enhancers of DRG hyperexcitability.** (A) Top ranked hit enhancers that increase the OA-SPARC phenotype. (B) Top ranked hit enhancers were validated using an 8-point, 3x dilutions series in both OA-SPARC sensitized- and uninflamed DRG neurons. The y-axis on the left indicates the phenotype score between the compound well and the average Vehicle (no SPARC) well. Shown here are four top-ranked hit enhancers, all of which enhanced the phenotype in a dose-dependent manner in OA-SPARC sensitized DRG neurons (purple). These compounds also induced an OA-like phenotype in uninflamed DRG neurons (yellow).

Supplement Table 1 | Validation of top ranked hits in rDRGs and rHippo

| Compound Name | MOA | Minimum distance in rDRG | Ralative total spikes in rHippo at minimum distance conc |
|---|---|---|---|
| Dibucaine | Na channel inhibitor | 1.05 | 1.33 |
| Nicardipine | Na channel inhibitor | 0.74 | 1.03 |
| FPL-64176 | Ion Channel Ligand | 0.84 | 0.79 |
| Gingerol | Ion Channel Ligand | 0.62 | 0.93 |
| Fipronil | Ion Channel Ligand | 0.54 | 0.81 |
| AZD6244 | MEK/MAPK signaling | 0.62 | 0.95 |
| Trametinib | MEK/MAPK signaling | 0.42 | 0.96 |
| Pimasertib | MEK/MAPK signaling | 0.33 | 0.84 |
| MEK162 | MEK/MAPK signaling | 0.37 | 1.15 |
| VRT752271 | MEK/MAPK signaling | 0.36 | 0.74 |
| Cobimetinib | MEK/MAPK signaling | 0.29 | 0.86 |
| S-Ruxolitinib | Tyrosine kinases modulator | 0.54 | 0.75 |
| Sunitinib | Tyrosine kinases modulator | 0.76 | 0.90 |
| Genistein | Tyrosine kinases modulator | 0.41 | 0.51 |
| Dovitinib | Tyrosine kinases modulator | 0.69 | 1.05 |
| Nintedanib | Tyrosine kinases modulator | 0.45 | 1.00 |
| Benidipine hydrochloride | Tyrosine kinases modulator | 0.54 | 0.35 |
| Romidepsin | HDAC inhibitor | 0.86 | N/A |
| Chidamide analog | HDAC inhibitor | 0.68 | 1.20 |
| Docusate sodium | GPCR | 0.83 | 0.62 |
| Ticagrelor | GPCR | 0.53 | 1.18 |
| Asenapine Maleate | GPCR | 0.36 | 0.22 |
| Tegaserod maleate | GPCR | 0.71 | 0.66 |
| AT13387 | Other mechanism | 0.62 | 1.13 |
| Licofelone | Other mechanism | 0.83 | 0.16 |
| Ro 31-8220 | Other mechanism | 0.46 | 0.55 |
| Fosaprepitant dimeglumine salt | Other mechanism | 0.91 | 0.53 |
| LY2784544 | Other mechanism | 0.49 | 0.76 |
| Aprepitant | Other mechanism | 0.66 | 0.46 |
| Cypermethrin | Other mechanism | 1.21 | 0.48 |
| Tubercidin | Other mechanism | 0.45 | 0.73 |
| Tiratricol | Other mechanism | 0.67 | 0.96 |
| Saracatinib | Other mechanism | 0.52 | 1.02 |

**Supplementary Table 1 | OA-SPARC reversal hits.** Top ranked hits were validated using an 8-point, 3x dilution series in both OA-SPARC sensitized DRG neurons and cultured rat hippocampal neurons. Minimum distance in OA-SPARC sensitized DRG neurons indicates efficacy. The total spikes relative to the vehicle (no SPARC) wells in rat hippocampal neurons at the minimal distance dose indicates the off-target effect in central neurons.
